# Dissecting the host determinants of flavivirus infection using QIC-seq

**DOI:** 10.1101/2025.07.15.664960

**Authors:** Allison J Dupzyk, Benjamin S Waldman, James Zengel, Jan E. Carette

**Affiliations:** Department of Microbiology and Immunology, Stanford University School of Medicine, Stanford, California, 94305, USA

## Abstract

Flaviviruses are genetically related, yet cause distinct disease patterns ranging from hepatitis and vascular shock syndrome to encephalitis and congenital abnormalities. There is an incomplete understanding of the cellular pathways co-opted by flaviviruses, and differences in host response to infection may underlie the diverse pathologies caused. We present a single-cell approach (Quantification of Infection and CRISPR guide sequencing; QIC-seq) that combines CRISPR/Cas9 knockout with virus-inclusive transcriptomics to systematically compare host factor requirements and host transcriptional response to flaviviral challenge. We show that dengue and yellow fever viruses are strictly dependent on subunits of the oligosaccharyltransferase complex, while the more distantly related West Nile and Langat viruses are dependent on components of the ER-associated degradation machinery. Our data further shows virus-induced upregulation of interferon-stimulated genes, and activation of the unfolded protein response. Together, QIC-seq enables quantitative comparisons of viral host factor utilization, which may inform development of host-directed antiviral therapies.

## Introduction

Flaviviruses are a genetically diverse genus of viruses, which pose a significant burden to global public health infecting more than 400 million people annually^1,2^. Dengue viruses (DENV) account for the majority of infections. The flavivirus genus includes a large group of tick-borne flaviviruses, which are prevalent in Europe and Asia, and a genetically distinct, large group of mosquito-borne flaviviruses, which are prevalent in tropical regions in Asia, Africa and South America^2–5^. Their geographic distribution will likely expand with climate change^6^. For several mosquito-borne flaviviruses including DENV, yellow fever virus (YFV) and Zika virus (ZIKV) humans and non-human primates are the primary host, whereas for other mosquito-borne flaviviruses including West Nile virus (WNV) and Japanese encephalitis virus, birds are the primary host. In addition to these variations in transmission cycle mediated by distinct insect vectors and mammalian hosts, flaviviruses also vary in their human pathology. Severe cases of WNV can cause neurotropic pathologies such as encephalitis, ZIKV infection can cause congenital disorders including microcephaly, and severe DENV and YFV infections can result in hemorrhagic fever.

Despite differences in their transmission cycle and pathologies, their replication strategy shares commonalities. Flaviviruses are enveloped, positive-sense, single-stranded RNA viruses with a genome approximately 11 kb in length^7^. The viral RNA, similar to messenger RNA, contains a 5′ cap, and encodes 10 viral proteins including 3 structural proteins, and 7 nonstructural proteins that together compose a large viral polyprotein. Flaviviruses are among a small group of positive-sense RNA viruses that lack a poly-A tail. Cotranslational insertion of the polyprotein into the endoplasmic reticulum (ER) membrane, and cleavage by host and viral proteases result in significant membrane reorganization to create invaginated viral replication organelles^8^. These ER membrane-localized replication organelles are a hallmark of flavivirus replication, and require polyprotein biogenesis and insertion.

High throughput CRISPR/Cas9, proteomic and siRNA screens have revealed that ER-resident host factors involved in proteostasis and quality control (QC) are essential for flavivirus infection^9–15^. Given the genetic diversity within the flaviviruses, it is unknown whether they have evolved common strategies to co-opt cellular pathways to promote their replication or whether there are differences between their host factor dependencies. Several of these host factors are transcriptionally upregulated during the unfolded protein response (UPR), an ER stress response which is activated during DENV infection^16–18^. This suggests that flaviviruses both induce the UPR and co-opt these factors to promote their replication.

Here, we developed an experimental strategy to systematically compare host factor requirements among divergent flaviviruses, and simultaneously capture the host transcriptomic response. This single-cell approach, which we termed **Q**uantification of **I**nfection and **C**RISPR guide sequencing (QIC-seq), combines CRISPR/Cas9 knockout with virus-inclusive transcriptomics. Our results functionally validate a core set of host factors involved in ER-proteostasis and quantify their contributions to RNA replication of four distinct flaviviruses. We identify cellular pathways induced by all tested flaviviruses, as well as specific genes upregulated only during challenge with specific flaviviruses. QIC-seq provides a much-needed protocol to systematically compare this diverse family of medically important viruses.

## Results

For development of QIC-seq (Fig. 1A) we chose to target cellular factors that in prior genome-scale screens were identified as having a role in DENV RNA replication, including those involved in ER-associated protein degradation (ERAD), N-linked glycosylation, and translocation and protein biogenesis^11,12,15,19^. Knockout cell lines of each host gene were generated by transduction of lentiviruses encoding single guide RNAs (sgRNAs) into Huh7.5.1 cells. Human hepatoma Huh7.5 and derivate cell lines including Huh7.5.1 are often used in studying flavivirus infection because they support robust replication and the liver is a target site for infection in human patients^20,21^. In total, 20 unique knockout cell lines were generated, each with a sgRNA targeting either a gene encoding a host factor previously identified as important in DENV infection, or a non-targeting (NT) guide as control. Amplicon sequencing of the targeted locus revealed efficient gene editing with insertion-deletion (indel) frequencies ranging between 80% - 95% for all sgRNAs except for the sgRNA targeting SND1, which showed less than 10% (Fig. 1B). Individual cell lines were pooled to obtain an equal ratio, and the resulting Huh7.5.1 cell library was then either mock infected, or infected with DENV type 2, for 48 hours.

**Figure 1:**
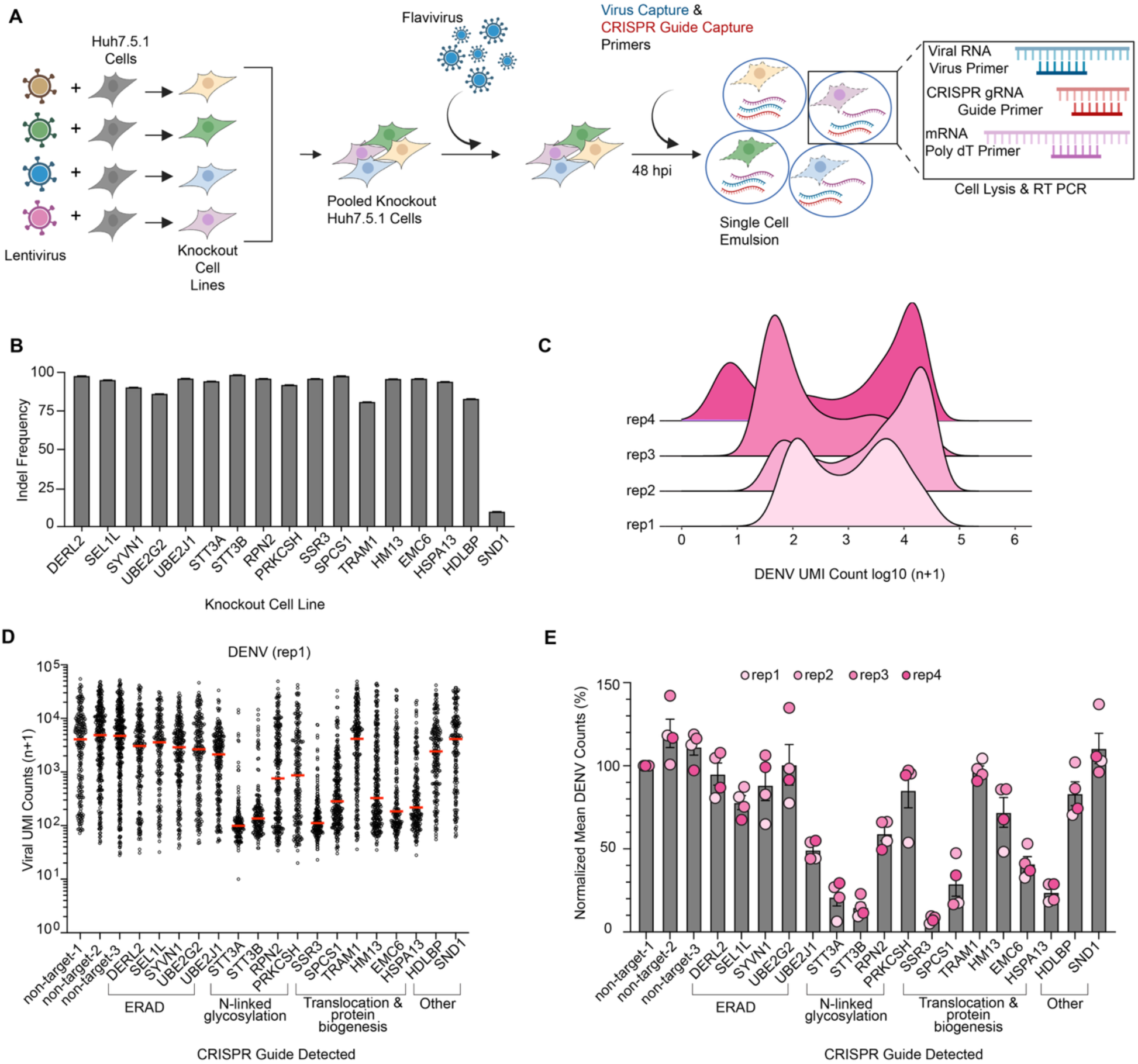
QIC-seq allows for intraviral comparison of host-factor requirements for DENV replication. **A,** Protocol design. Individual lentiCRISPRv2 plasmids were used to generate knockout cell lines that were pooled to generate a cell library. Cells were unchallenged, or challenged with DENV for 48 hours, harvested, and subject to 5′ capture using the 10X Genomics 5′ VDJ kit, with addition of primers targeting the 5′ region of the viral RNA, or the CRISPR guide scaffold. **B,** Indel frequency of Huh7.5.1 knockout cell lines. **C**, DENV UMI counts (log_10_ of n+1) in all guide detected, DENV challenged Huh7.5.1 cells, separated by each biological replicate of DENV QIC-seq screen. **D**, QIC-seq cell plot of first biological replicate of DENV QIC-seq screens. Cells are plotted as circles by guide-detected and DENV UMI counts (log_10_ of n+1). Red line represents the median value. **E,** Graph of the mean DENV count in guide detected cells, divided by the mean DENV count in non-target-1 guide detected cells for each biological replicate. Four biological replicates are graphed. Error bars represent standard error of the mean.

Cells were collected and prepared for single cell RNA sequencing using the 10X 5′ VDJ platform. To allow for the amplification of CRISPR guide RNA and DENV genomic RNA, we added capture primers annealing to the conserved loop in the sgRNA and to the 5′-end of DENV, respectively^22^. This modification in the protocol ensures that, like the polyadenylated mRNA, sgRNAs and DENV genomic RNA cDNAs incorporate a unique modular identifier (UMI) to limit the impact of PCR bias in later steps, and a cell barcode which allows the assignment of the sgRNA, viral RNA and cellular mRNA to single cells and enables simultaneous evaluations of viral RNA replication with identification of host perturbations (Fig. 1A). After reverse transcription and cDNA amplification, cDNAs encoding the 5′ end of the viral genome and the sgRNA were size separated from cDNAs derived from polyadenylated mRNA (transcriptome) before processing both libraries for single cell sequencing. Alignment generates a matrix of mRNA expression values for each cell, as well as a table with viral UMI counts and CRISPR guide identification for each cell, enabling quantification of the effects of CRISPR perturbation on viral replication, and insight into the host transcriptomic response.

### DENV replication is quantifiable in guide-detected cells

Following quality control thresholding and elimination of cells with greater than 1 guide identified, we analyzed DENV counts in 11,114 DENV challenged Huh7.5.1 cells across 4 biological replicates. As a control, we similarly analyzed a cell population that was not infected by DENV (unchallenged); in these 5,070 cells, there were no detectable DENV UMI counts (n + 1 = 1) in > 99% of cells (Supplemental Table 1). DENV UMI counts in DENV challenged Huh7.5.1 cells display a generally bimodal distribution, likely representing two populations of cells-one population of infected cells where replication of DENV RNA is occurring, and one non-infected population with minor to no DENV replication (Fig. 1C). To directly compare the UMI counts of the lower peak to uninfected cells, we spiked in unchallenged cells just prior to droplet generation (Supplemental Fig. 1A). We found this population displayed low, but measurable, DENV UMI counts, and this distribution is comparable to that of the lower peak in infected samples, reinforcing the notion that these cells have not supported appreciable DENV RNA replication (Supplemental Fig. 1B).

Separating cells based on the identified sgRNA enables comparison of how individual host factor knockouts impact DENV viral replication. In these graphs, which we term QIC-seq plots, individual cells are plotted based on DENV UMI count and split by guide detected (Fig. 1D). The majority of non-target guide detected cells have DENV UMI counts greater than 1,000. In contrast, the large majority of cells lacking host factors previously demonstrated to be required for DENV replication, such as the catalytic subunits of the oligosaccharyltransferase (OST) complex, STT3A and STT3B, show an approximately 100-fold reduction in DENV UMIs, comparable to the reduction observed in our spike in experiment and indicating a lack of replication. A similarly strong reduction in DENV replication was observed in cells with guides targeting SSR3, EMC6 and HSPA13, while targeting UBE2J1, RPN2, and PRKCSH resulted in more moderate reductions. Although the median DENV count in non-target guide detected cells varies between replicates, the overall trend remains the same, highlighting the reproducibility of this screening method (Fig. 1E, Supplemental Fig. 1C-E). Thus, we have established QIC-seq as an experimental strategy that allows for quantitative intraviral comparisons of host factor requirements.

### Divergent flaviviruses have dependencies on unique host factors

Mosquito-borne flaviviruses such as DENV, WNV and YFV comprise a clade within the flaviviruses that is distinct from the clade of tick-borne viruses such as LGTV^23^. The mosquito-borne flaviviruses can be divided into those that use birds as reservoirs such as WNV, and those that use primates as their primary mammalian host such as DENV and YFV. Our library of ER-proteostasis host factors was based on extensive screening with DENV, however, the conservation of these host factor requirements across diverse members of the family has not been well-characterized. We applied QIC-seq to determine the effect of knockout of host factors on viral replication and host response. Our Huh7.5.1 cell library was challenged with LGTV, WNV or YFV for 48 hours, and again subject to capture and QIC-seq library preparation. The multi-virus Huh7.5.1 QIC-seq screen contained 1,671, 2,171, and 3,602 LGTV, WNV, and YFV challenged cells, respectively, which we merged with the initial 11,114 DENV challenged and 5,070 unchallenged cells (Supplemental Table 1) to compare host transcriptional response and viral replication in perturbation identified cells. Analysis of the viral UMI counts again revealed a bimodal distribution (Supplemental Fig. 2A).

Generally, host factors important for DENV were also important for other flaviviruses, likely reflecting a conserved role of these cellular factors in flavivirus polyprotein biogenesis required for RNA replication (Fig 2A-2C). However, we did observe intriguing differences. While knockout of STT3A and STT3B results in strong decreases in DENV and YFV RNA replication, the other flaviviruses were not or only slightly affected (Fig. 2D). Conversely, knockout of ERAD components affected LGTV and WNV RNA replication, had a moderate effect on DENV, but had little to no effect on YFV replication (Fig. 2D). Knockout of the translocon associated protein SSR3 and EMC6, part of 2 protein complexes required for insertion of transmembrane domains at the ER^24^, led to strong decreases in viral replication for all flaviviruses with the notable exception of SSR3 for WNV (Fig. 2B, 2D).

**Figure 2:**
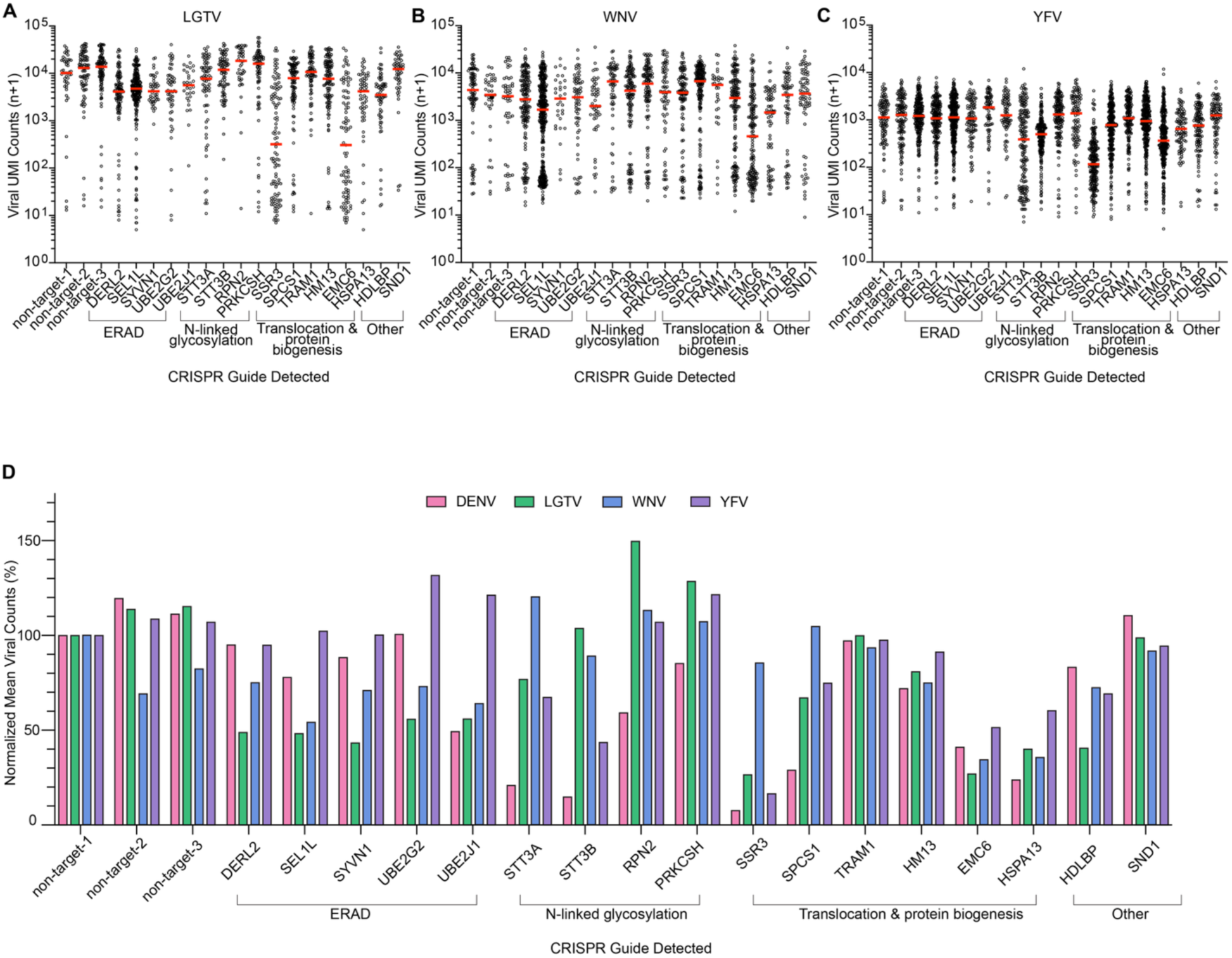
Divergent flaviviruses have unique host factor dependencies. **A**, QIC-seq plot of LGTV challenged Huh7.5.1 cells. Cells, represented as circles, are plotted by guide detected and LGTV UMI counts (log_10_ of n+1). Red line represents median value. **B,** As in **A** except cells are challenged with WNV. **C**, As in **A** except cells are challenged with YFV. **D**, Mean viral UMI counts in guide detected cells, normalized to the mean viral counts of non-target-1 guide detected cells in each viral challenge.

### Universal upregulation of the unfolded protein response by divergent flaviviruses

QIC-seq combines CRISPR perturbation with a transcriptional readout of cellular mRNA during viral infection. After dataset integration and normalization, we first compared the RNA profiles of cells present in the DENV challenged samples to cells present in the unchallenged sample. Gene ontology revealed the UPR and ERAD as the most enriched categories (Supplemental Fig. 2B). This is in line with prior reports indicating that UPR activation occurs during flavivirus infection^25^. No obvious activation of interferon stimulated genes was observed. To facilitate further comparative analysis, we assembled a list of genes that have been experimentally shown to be transcriptionally upregulated during ER stress (UPR module)^26,27^ and a list of interferon stimulated genes (IFN module)^28,29^ (Supplemental Table 2). Of the 113 genes upregulated in the DENV challenged cells, 48 genes corresponded to the UPR master list (42%), and 2 genes corresponded to the IFN stimulated master list (2%) (Fig. 3a, Supplemental Table 3). Dimensional reduction analysis was used to visualize DENV UMI counts in individual cells (Fig. 3B, left plot). As a quantitative measure of the gene set activity of different pathways, we calculated module scores for our UPR and IFN gene lists. We found that cells with high DENV-UMI reads also displayed high UPR module scores, while no such relationship was found for the IFN module (Fig. 3B, center and right plot).

**Figure 3:**
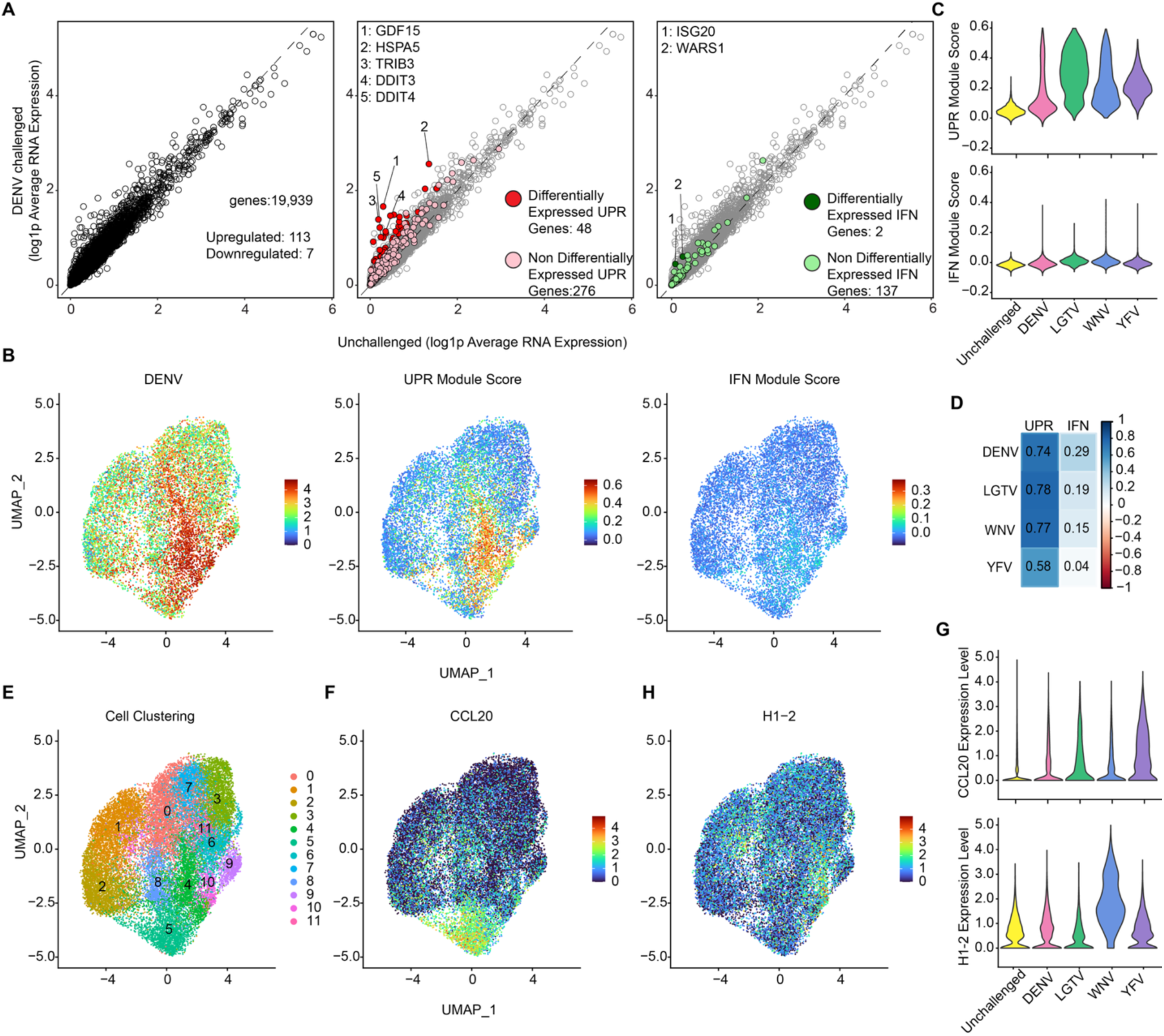
Divergent flaviviruses collectively induce the UPR in Huh7.5.1 cells. **A**, Log1p average RNA expression of genes in DENV challenged and unchallenged cells. Circles represent genes, y=x line denotes genes equally expressed between both challenged and unchallenged cells. Left graph: all genes detected in cells in black. Center graph: All genes not corresponding to the UPR master list: light grey. Differentially expressed genes corresponding to UPR master list: red. Non-differentially expressed genes corresponding to UPR master list: pink. Right graph: All genes not corresponding to IFN stimulated gene master list: light grey. Differentially expressed genes corresponding to IFN stimulated gene master list: dark green. Non-differentially expressed genes corresponding to IFN stimulated gene master list: light green. **B**, UMAP visualization of DENV challenged Huh7.5.1 cells. Features include DENV UMI counts (log_10_ n+1 ), UPR module score, and IFN stimulated gene module score. **C**, Violin plot of UPR and IFN stimulated gene module scores of all virally challenged and unchallenged Huh7.5.1 cells, separated by viral challenge. **D**, Correlation plot of viral counts and UPR and IFN stimulated gene module score in Huh7.5.1 cells. Spearman’s correlation coefficient used. **E**, UMAP of all Huh7.5.1 cells, clustered by gene expression. **F**, Feature plot of all Huh7.5.1 cells, with CCL20 expression featured. **G**, top: Violin plot of CCL20 gene expression, separated by viral challenge. Bottom: Violin plot of H1-2 gene expression, separated by viral challenge. **H**, Feature plot of all Huh7.5.1 cells, with H1-2 expression featured.

Plotting module scores of cells present in the DENV-challenged sample and comparing them to the unchallenged sample further indicates activation of UPR (Supplemental Fig. 2C, D). Within the DENV infected population, cells with non-target sgRNAs exhibited strong UPR activation, while cells containing sgRNAs targeting DENV-essential host factors such as STT3A and STT3B displayed much reduced activation (Supplemental Fig. 2D). This suggests that DENV RNA replication is the main driver of UPR activation, which can be prevented by knockout of virus-essential host factors. We did observe a modest increase in UPR module scores in unchallenged cells with guides targeting SEL1L, UBE2G2, STT3A and HSPA13 albeit to a much lesser extent than the UPR activation observed in virally challenged cells (Supplemental Fig. 2C). Analysis of cells challenged with LGTV, WNV and YFV displayed a similar increase of UPR module score but not IFN when compared to the unchallenged population (Fig. 3C). While cells do not appear to cluster based on type of viral challenge, cells with the highest viral counts in all challenges do cluster together (Supplemental Fig. 3a-B). For all tested flaviviruses, viral RNA levels correlated positively with the UPR module score, and again we observed cells with high viral UMI counts also displayed high UPR module scores (Fig. 3C-D, Supplemental Fig. 3B). This suggests that the upregulation is due to a cell-intrinsic mechanism and not paracrine signaling. Correlations for IFN were negligible.

We clustered cells using k-nearest neighbors to further explore differential gene expression (Fig. 3E, Supplemental Table 3). Cells in cluster 4 displayed strong upregulation of UPR genes, and also corresponded to cells with the highest viral counts for all viruses (Fig. 3B, 3E, Supplemental Fig. 3B). The highest upregulated genes in cells in cluster 5 included several chemokine ligands such as CCL20, CXCL1, and CXCL8. Cells in clusters 6 and 10 showed an upregulation of several histone genes including H2AC17, H2BC18, and H2AC13. Gene ontology analysis did not demonstrate an obvious upregulation of these pathways comparing unchallenged cells to virus-challenged cells, likely due to the magnitude of UPR upregulation during viral challenge (Supplemental Fig. 3C-E), though we observe upregulation of several individual chemokine ligand genes during infection, especially in YFV-challenged cells (Fig. 3F-G, Supplemental Table 3). Several histone genes appeared to be upregulated specifically during WNV infection (Fig. 3G-H, Supplemental Table 3). Thus, while divergent flaviviruses strongly induce a common UPR, they may also induce more specific transcriptional pathways.

### Optimized QIC-seq protocol for pooled library generation

Initial QIC-seq screens were performed by combining individually-generated knockout cells, allowing us to determine gene editing efficiency of the individual sgRNAs and ensure each cell received a single sgRNA. To optimize this protocol for single-step cell library generation, we generated a single-vector lentiviral CRISPR library targeting the same 17 genes with 4 guides per gene. In addition to four non-targeting control guides, we included guides targeting KIAA0319L, a gene not thought to be involved in flavivirus infection. This substantially quickens the cell library generation phase and streamlines the QIC-seq protocol (Fig. 4A). To compare the transcriptional responses to flavivirus infection and their dependencies on host factors between different cell types, we chose to introduce the library in HAP1 cells, which have been used in prior genetic screens for flaviviruses^11,30^. HAP1 cells were either challenged with virus, or unchallenged for 48 hours, and subject to QIC-seq library preparation.

**Figure 4:**
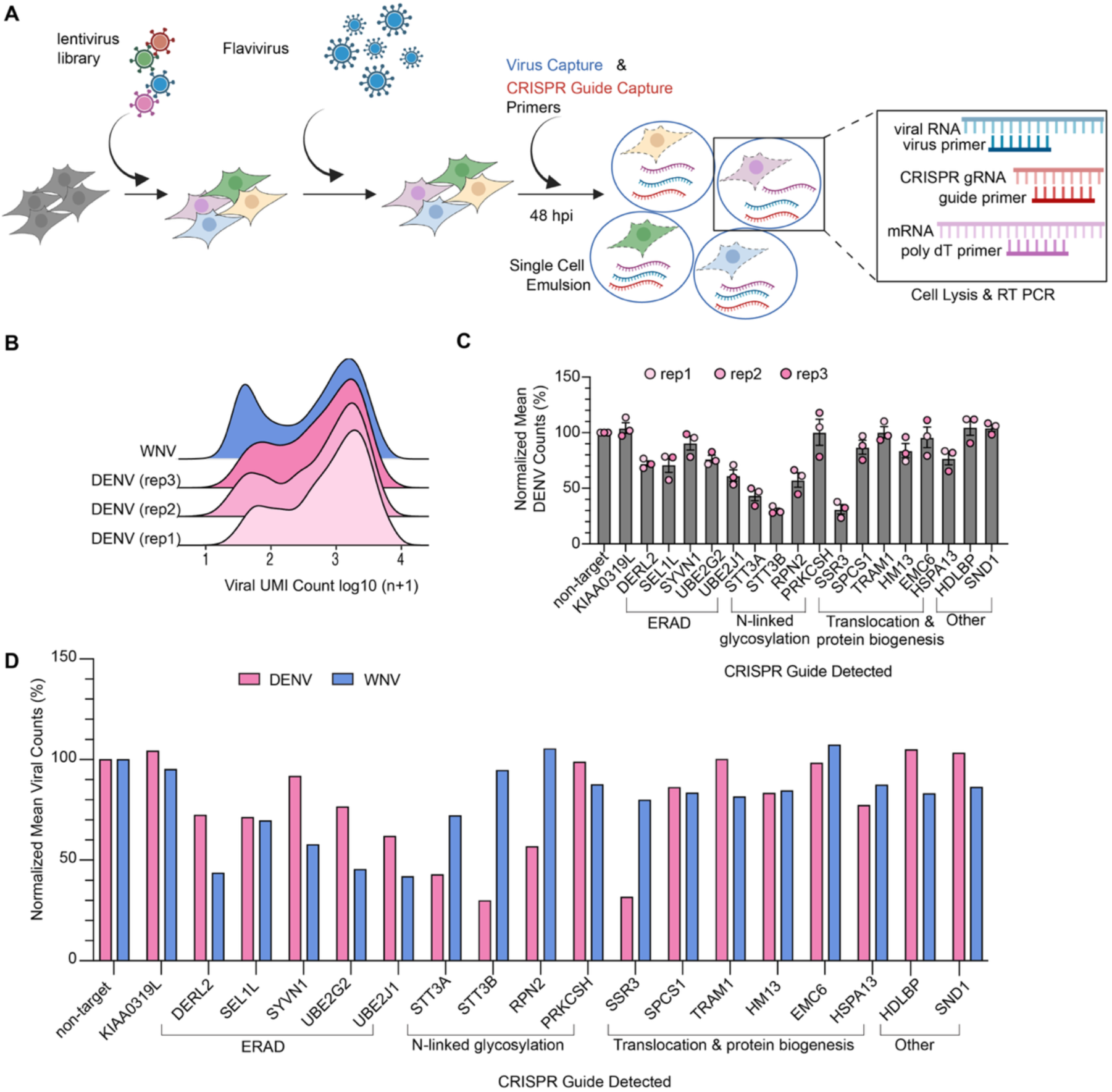
DENV replication is dependent on STT3A and STT3B in HAP1 cells, while WNV replication is dependent on ERAD machinery. **A**, Optimized QIC-seq protocol design. LentiCRISPRv2 plasmid library was used in single transduction to generate a cell library. Cells were unchallenged, or challenged with DENV or WNV for 48 hours, harvested, and subject to 5′ capture using the 10X Genomics 5′ VDJ kit, with addition of primers targeting the 5′ region of the viral RNA, or the CRISPR guide scaffold. **B**, UMI counts (log_10_ of n+1) in viral challenged HAP1 cells, separated by virus, and biological replicate. Replicate 1, 2 and 3 are challenged with DENV at an MOI of 5, 7, and 9, respectively. **C**, Mean DENV count in guide detected cells, divided by mean DENV count in non-target guide detected cells for each biological replicate. Three biological replicates are graphed with error bars representing standard error of the mean. **D**, Mean viral count in HAP1 guide detected cells, divided by the mean viral count in non-target guide detected cells for each biological replicate. Three biological replicates are included in the DENV challenge.

In total 2,972 unchallenged HAP1 cells were compared to 6,133 DENV challenged and 3,770 WNV challenged HAP1 cells. The DENV infection was performed in triplicate using an MOI of 5, 7, and 9, whereas the WNV infection was performed once at an MOI of 2.5. Like before, we observed a bimodal distribution of viral read counts representative of different degrees of RNA replication in individual cells (Fig. 4B, Supplemental Table 4). To analyze the phenotypic consequences of perturbation on flavivirus replication in HAP1 cells, we plotted cells by viral counts and guide detected using QIC-seq plots (Supplemental Fig. 4A). We observed consistent results between the three DENV replicates (Fig. 4C). Generally, dependency profiles were similar to those observed in Huh7.5.1 cells with knockout of STT3A, STT3B, and SSR3 showing the most pronounced decrease in viral RNA counts and the components of the ERAD pathway showing a more moderate reduction. However, knockout of HSPA13 had a much more moderate effect in HAP1 cells, and, most strikingly, EMC6 knockout did not affect RNA replication in HAP1 cells while in Huh7.5.1 cells it resulted in strong replication defects. Comparing DENV with WNV in HAP1 cells revealed a stronger dependence of WNV on ERAD components and no or only a slight dependence on STT3A and SSR3, which was similar to the results in Huh7.5.1 (Fig. 4D, Supplemental Fig. 4B). Like DENV, the strong dependence on EMC6 observed in Huh7.5.1 cells was not observed in HAP1 cells. Thus, we have optimized the QIC-seq experimental strategy and shown its utility in facilitating quantitative analysis of flavivirus replication.

### QIC-seq reveals transcriptional differences in HAP1 cells challenged by DENV and WNV

We next investigated the transcriptional responses upon viral challenge in HAP1 cells. Intriguingly, while UMI distributions of WNV and DENV were comparable (Fig. 4B), the transcriptional response to DENV infection was subdued compared to WNV with only 121 upregulated genes versus 902 in the WNV challenged cells (Fig. 5A-B, Supplemental Table 5). Gene ontology analysis of the DENV upregulated genes showed terms related to ribosome biogenesis and cytoplasmic translation but genes involved in the UPR or IFN response were not enriched (Supplemental Fig. 5A). In stark contrast, WNV challenged cells showed a strong statistical enrichment for gene sets involved in the response to ER stress (UPR) and the defense response to virus (IFN signaling) (Supplemental Fig. 5B). In line with this, cells in the WNV challenged population exhibited an upregulation of 102 genes from the UPR master list including widely used UPR markers such as DDIT3 (also named CHOP/GADD153) and HSPA5 (also named BiP/GPR78), and 57 genes from the IFN list including well-characterized interferon stimulated genes including ISG15 and IFITM1 (Fig. 5B, Supplemental Table 5). No such upregulation was observed for DENV challenged cells (Fig. 5A). Gene set activity analysis using module scores further underscored the pronounced activation of the UPR and IFN pathways in cells challenged with WNV, but not DENV (Fig. 5C). Within the DENV infected sample we observed little to no correlation between viral count numbers and the IFN or UPR module scores (Fig. 5D). As observed for the Huh7.5.1 cells, WNV read counts were positively correlated with the UPR module score. Interestingly, we found that the IFN module score was negatively correlated with WNV. Dimensional reduction analysis showed clear separation between unchallenged, DENV-challenged and WNV-challenged cells (Fig. 5E). Elevated UPR and IFN module scores were observed in the region corresponding to the WNV-challenged sample (Fig. 5F). Plotting the WNV counts revealed a clear separation in the UMAP between highly infected cells and lowly-infected “bystander” cells (Fig. 5F, right). The IFN module score was highest in the bystander cells suggesting that WNV infection triggers a paracrine interferon response that is suppressed by viral replication. As in Huh7.5.1 cells, we observed an upregulation of several core and linker histone proteins in the WNV-infected sample, but this upregulation was not obviously correlated to WNV counts (Fig. 5G). No chemokine ligand upregulation, as seen in some virus-challenged Huh 7.5.1 cells, was observed (Supplemental Fig. 5C-D). Together, these experiments demonstrate that WNV infection triggers the UPR and IFN response in HAP1 cells, and that viral counts positively correlate with UPR activation and negatively with IFN response.

**Figure 5:**
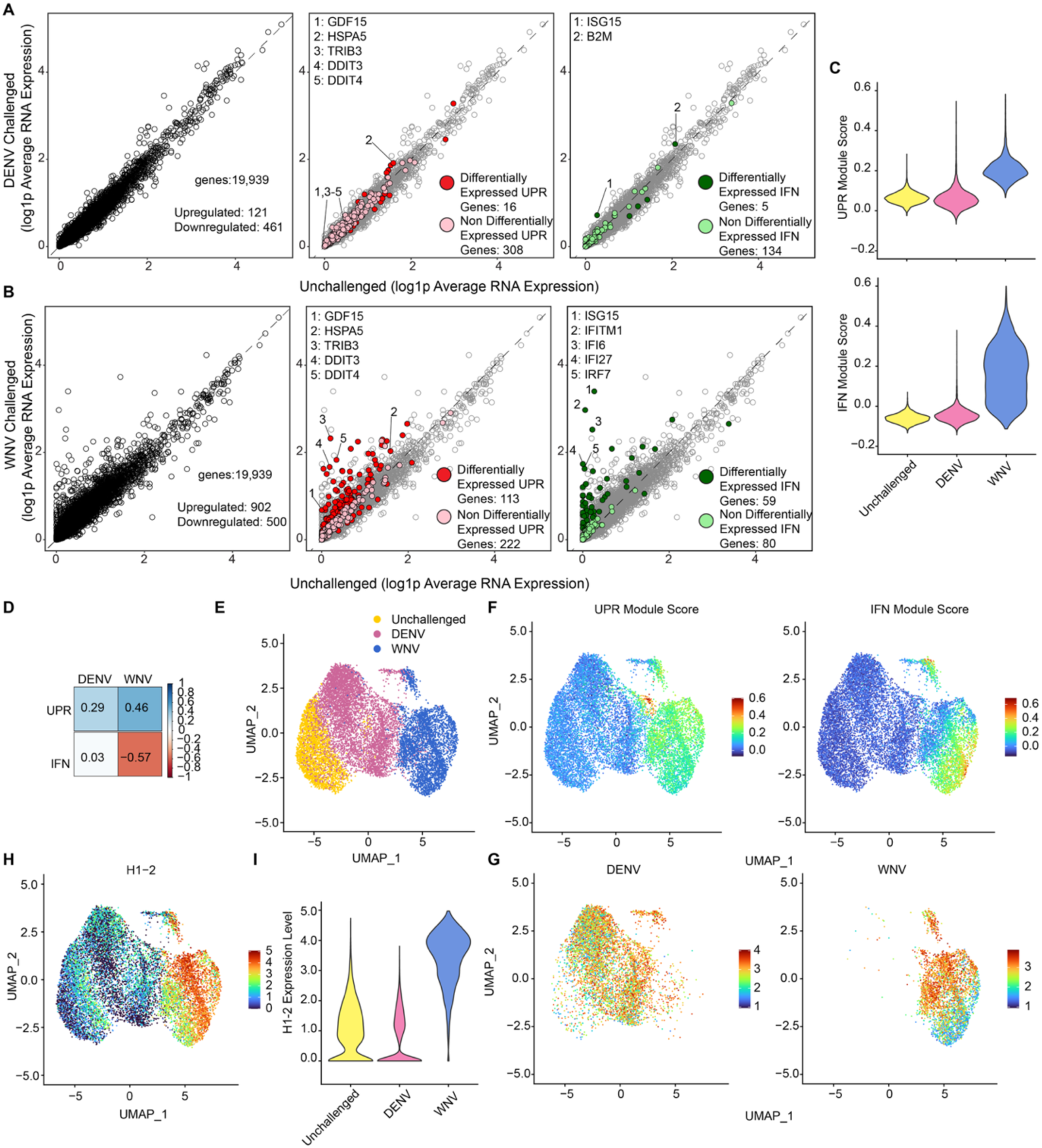
WNV challenge induces UPR and an IFN response in HAP1 cells. **A**, Log1p Average RNA expression of genes in DENV challenged and unchallenged cells. Circles represent genes, y=x line denotes genes equally expressed between both challenged and unchallenged cells. Left graph: all genes detected in cells in black. Center graph: All genes not corresponding to the UPR master list: light grey. Differentially expressed genes corresponding to UPR master list: red. Non-differentially expressed genes corresponding to UPR master list: pink. Right graph: All genes not corresponding to IFN stimulated master list: light grey. Differentially expressed genes corresponding to IFN stimulated master list: dark green. Non-differentially expressed genes corresponding to IFN stimulated master list: light green. **A**, As in Figure **A**, however, cells are challenged with WNV. **C**, Violin plots of UPR and IFN stimulated gene module scores in HAP1 cells, separated by virus challenge. **D,** Correlation plot of viral counts and UPR and IFN stimulated gene module score in HAP1 cells. Spearman’s correlation coefficient used. **E**, Feature plot of all HAP1 cells, colored by challenge. **F**, Feature plots of HAP1 cells. Features include: top left: UPR module score, top right: IFN module score of all HAP1 cells, bottom left: DENV UMI counts (log_10_ n+1 ) in DENV challenged HAP1 cells and bottom right: WNV UMI counts (log_10_ n+1 ) in WNV challenged Hap1 cells. **G**, Left: Feature plot of H1-2 expression in HAP1 cells. Right: Violin plot of H1-2 expression in HAP1 cells, separated by viral challenge.

## Discussion

We have developed a single-cell RNA sequencing strategy (QIC-seq) that allows for quantification of non-polyadenylated RNA virus replication, perturbation identification, and single cell transcriptional analysis. Conventional CRISPR fitness screening approaches rely on bulk PCR amplification of sgRNA sequences from genomic DNA of cell populations to infer importance for viral fitness based on survival of viral challenge. In contrast, QIC-seq is an RNA-based approach that directly and sensitively measures the impact of host factor knockout on viral RNA accumulation and mRNA abundance. A recent study independently developed a similar approach to analyze SARS-CoV-2 replication and the interferon response in cells depleted of host factors identified as viral-binding proteins^31–35^. This approach has similarities with QIC-seq but is not suitable for non-polyadenylated viruses such as flaviviruses.

RNA virus-inclusive single-cell RNA-sequencing has been instrumental in multiple studies monitoring heterogeneity in host response to viral challenge^31–35^. We have used QIC-seq to systematically compare host factor dependencies and transcriptional responses between divergent flaviviruses and between distinct cell types. Notably, QIC-seq enables the monitoring of host transcriptional responses under viral challenge and amidst perturbation. These responses include innate immune responses. Because host-directed antiviral development involves both direct targeting of host dependency factors, but also modulating immune responses, QIC-seq could provide a versatile tool to be used in this context^36^.

Our QIC-seq plots revealed unique host factor requirements for flaviviral replication. DENV RNA accumulation was consistently reduced in cells where STT3A or STT3B guides were detected to levels comparable to those observed in cells not exposed to DENV. This strict DENV-dependence on the OST catalytic subunits is in line with prior reports^11,12^. The same observation was true of YFV replication. In contrast, WNV and LGTV showed little to no dependency on either STT3A or STT3B and instead depended more strongly on the core components of the ERAD machinery. This suggests that divergent flaviviruses have evolved distinct mechanisms to utilize these host factors, although further studies are needed to reveal the molecular details.

Flavivirus infection induces ER stress due to the high expression level and complex topology of the viral polyprotein. The strong positive correlation observed between viral UMI counts and the UPR module score in virus-infected cells demonstrates QIC-seq’s ability to identify cellular pathways induced during viral infection. The role of the UPR in promoting or antagonizing viral replication remains enigmatic^16^. While not undertaken here, QIC-seq is a useful tool that could be used to further our understanding of the UPR-flavivirus relationship by generating and testing a CRISPR library specifically focused on components of the UPR pathway.

We also observed a small number of genes upregulated in cells challenged with specific viruses. For example, the histone-encoding gene H2AC13 is the second most upregulated gene in WNV-challenged Huh7.5.1 cells when compared to unchallenged cells. In both Huh7.5.1 and HAP1 cells, the histone gene H1-2 is also highly upregulated during viral challenge.The DENV capsid protein has been shown to bind to core histones, inhibit nucleosome formation, and increases the protein expression levels of core histones^37^. Our results with WNV suggest that histone upregulation is more broadly seen during flavivirus infection and occurs at the transcriptional level. The transcriptional pathway leading to this upregulation and whether this stimulates or antagonizes viral infection is unclear. We also observed differential expression of chemokine ligands, such as CCL20, in a small group of Huh7.5.1 cells, which may be explained by ER stress leading to inflammation through CREBH and ATF6 cleavage^38^.

QIC-seq screens in HAP1 cells allowed us to investigate the IFN response to flaviviruses, as these cells have intact RIG-I signaling, unlike Huh7.5.1^29,39,40^. In WNV-challenged HAP1 cells, we found a strong activation of IFN stimulated genes suggesting that WNV infection results in the production of interferon. Intriguingly, we observed a negative correlation between WNV UMI counts and the IFN module score, likely indicative of paracrine IFN signaling, which is antagonized in highly infected cells. This demonstrates the ability of virus-inclusive scRNA-seq to uncover complex relationships between viral infection and transcriptional responses, as bulk RNA-sequencing experiments would have masked the inverse relationship in WNV-infected cells. WNV nonstructural proteins have been shown to inhibit IFN signaling through preventing phosphorylation of STAT1 and STAT2^41,42^.

Interestingly, despite DENV UMI counts indicating productive infection in HAP1 cells, no correlation with the IFN module score was observed. This suggests that DENV infection did not result in production of IFN. This could be due to the superiority of DENV in preventing the initial step in the production of IFN, which is the detection of viral pathogen-associated molecular patterns (PAMPs) by pattern recognition receptors (PRRs) such as RIG-I and cGAS. Whether this reflects a passive mechanism by hiding PAMPs^43,44^ or active suppression by inactivating PRRs^45^ is unknown. Given the minimal UPR activation in DENV challenged HAP1 cells, it is also possible that lower efficiency of RNA replication compared to WNV infection contributed to the failure to trigger an IFN response.

In summary we present QIC-seq, a single-cell RNA-sequencing method in which flavivirus replication is paired with perturbation identification and host transcriptomics. Using QIC-seq, we have compared divergent flavivirus dependence on two major proteostasis-maintaining host complexes, the OST complex and ERAD machinery, and find unique host factor requirements for replication. We have identified the UPR as commonly upregulated during flavivirus challenge, and identified small numbers of unique histone and chemokine ligand genes uniquely upregulated during viral challenges. This work furthers our knowledge of cellular requirements of a group of medically important viruses that cause significant burdens to global public health, with the potential to inform therapeutic design.

---

**Supplemental Figure 1:**
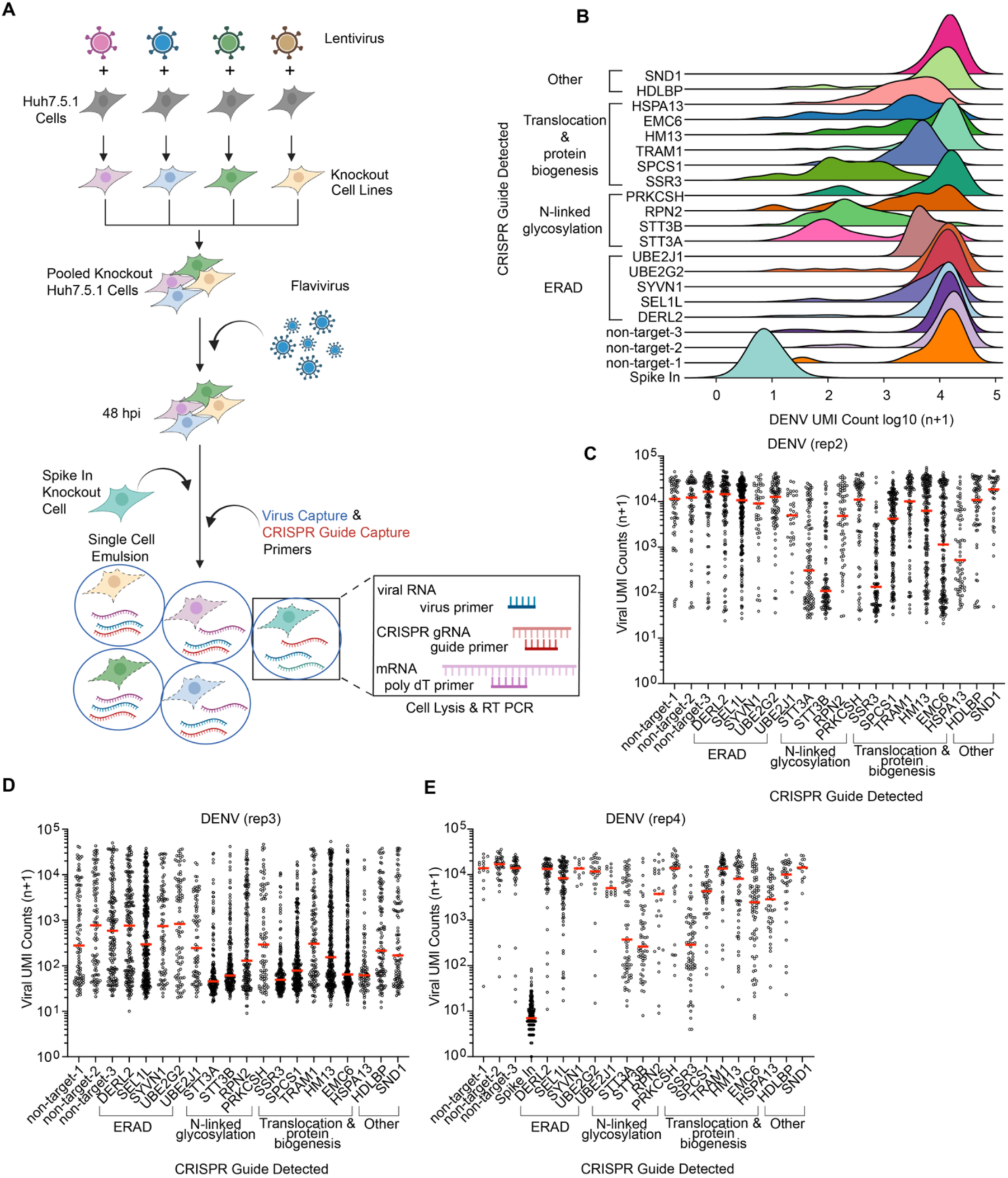
Replicate QIC-seq screens quantify the effect of knockout of select host genes on DENV RNA replication. **A-C**, QIC-seq cell plot of remaining three biological replicates of DENV QIC-seq screen. Cells, represented as circles, are plotted by guide detected and DENV UMI counts (log_10_ of n+1). Red line represents median value. **A**, Protocol design for spike in experiment. As in **1A**, however, naïve knockout Huh7.5.1 cells are added to the challenged cell library just before single cell emulsion. These “Spike In” cells are identified by the expression of a unique sgRNA. **B**, Ridgeplot of spike in experiment showing the distribution of DENV UMI counts (log_10_ of n+1). **C,** Second biological replicate of DENV QIC-seq screen. **D**, Third biological replicate of DENV QIC-seq screen. **E,** Fourth biological replicate of DENV QIC-seq screen (spike in experiment). This replicate has a slight modification to the protocol as outlined in subpanel **A**.

**Supplemental Figure 2:**
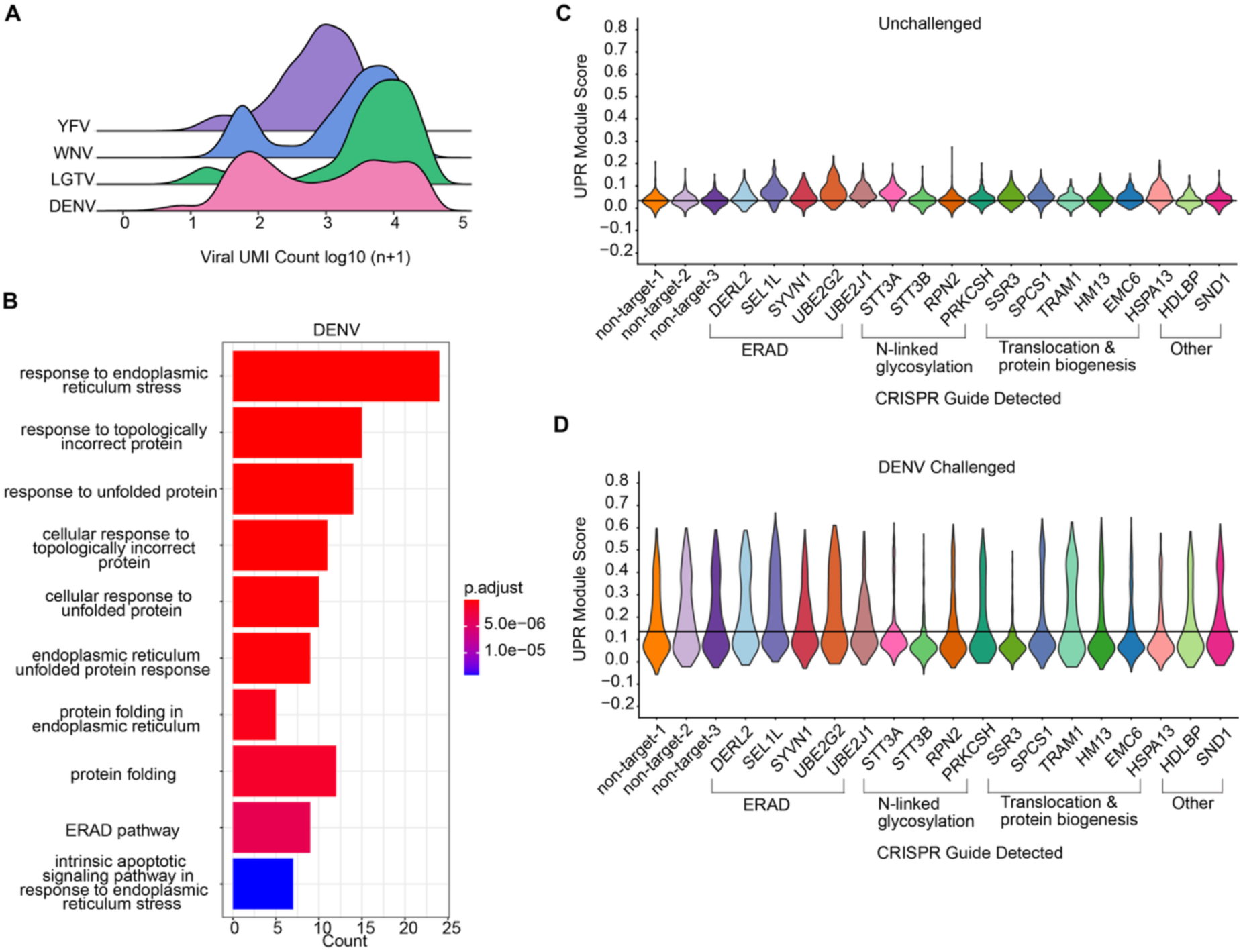
DENV induces the UPR in Huh7.5.1 cells. **A**, UMI counts (log10 of n+1) in viral challenged Huh7.5.1 cells, separated by virus. DENV ridgeplot includes all four biological replicates. **B**, GO plot of pathways upregulated based on DEGs found between unchallenged and DENV challenged Huh7.5.1 cells. List of genes generated using log2fc 20.25. Top 10 most upregulated pathways plotted. Top 10 most upregulated pathways plotted. **C**, Violin plot of UPR module scores in unchallenged Huh7.5.1 cells, grouped by guide detected. Line represents median value of module scores in non-target guide detected cells. D, As in supplemental **C**, however cells have been challenged with DENV.

**Supplemental Figure 3:**
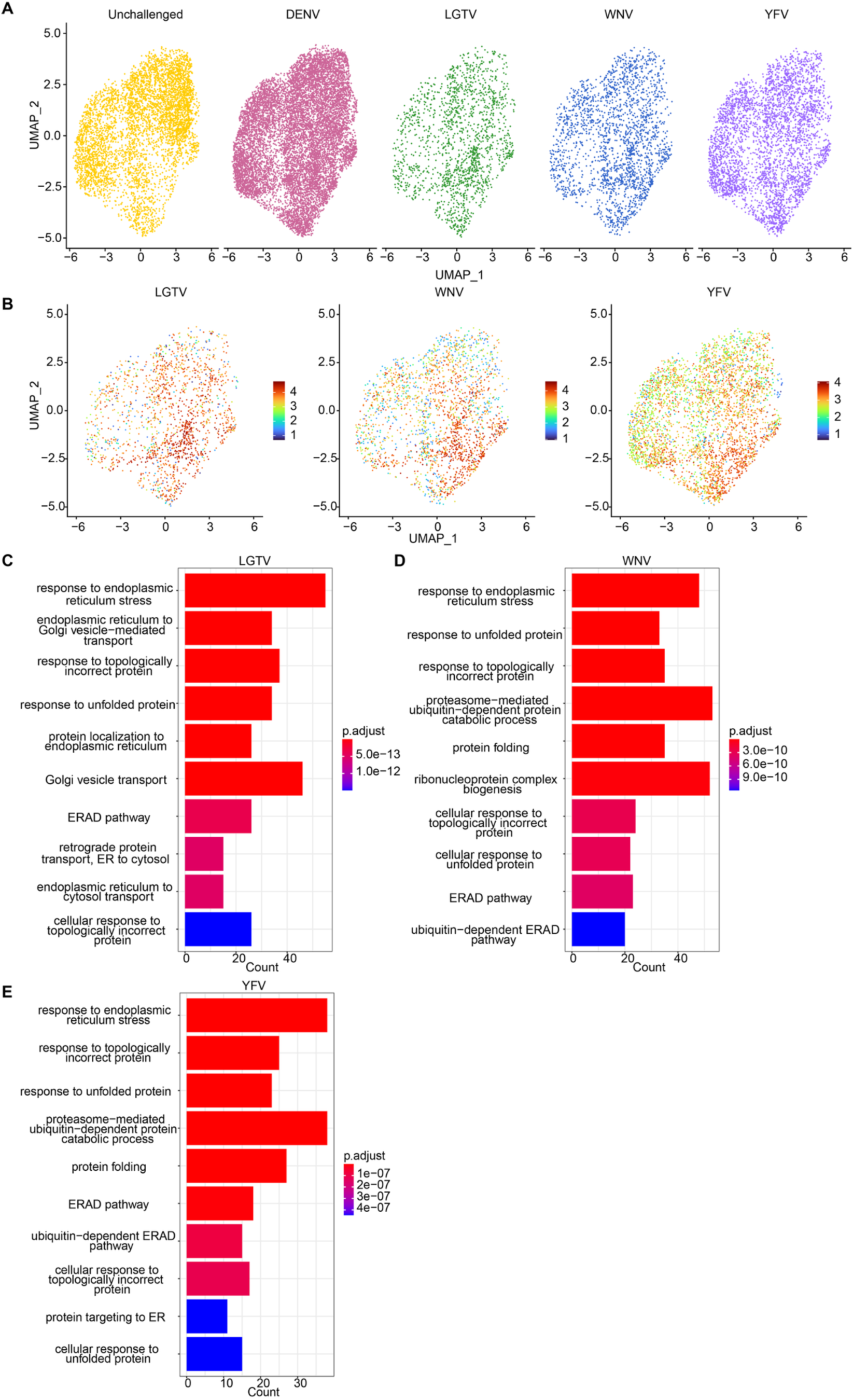
UMAPs and gene ontology analysis of Huh7.5.1 cells challenged with divergent flaviviruses. **A**, UMAP of Huh7.5.1 cells, split and colored by viral challenge. Only Huh7.5.1 cells challenged by indicated virus, or unchallenged cells are displayed for each plot. **B**, Feature plot of Huh7.5.1. Features include LGTV, WNV, and YFV UMI counts (log10 n+1). Only Huh7.5.1 cells challenged by indicated virus are displayed for each plot. **C**, GO plot of pathways upregulated based on DEGs found between LGTV challenged and unchallenged Huh7.5.1 cells. List of genes generated using log2fc ≥ 0.25. Top 10 most upregulated pathways plotted. Count represents the number of DEGs belonging to the indicated pathway. **D**, As in **C**, however, cells are challenged with WNV. **E**, As in **C**, however cells are challenged with YFV.

**Supplemental Figure 4:**
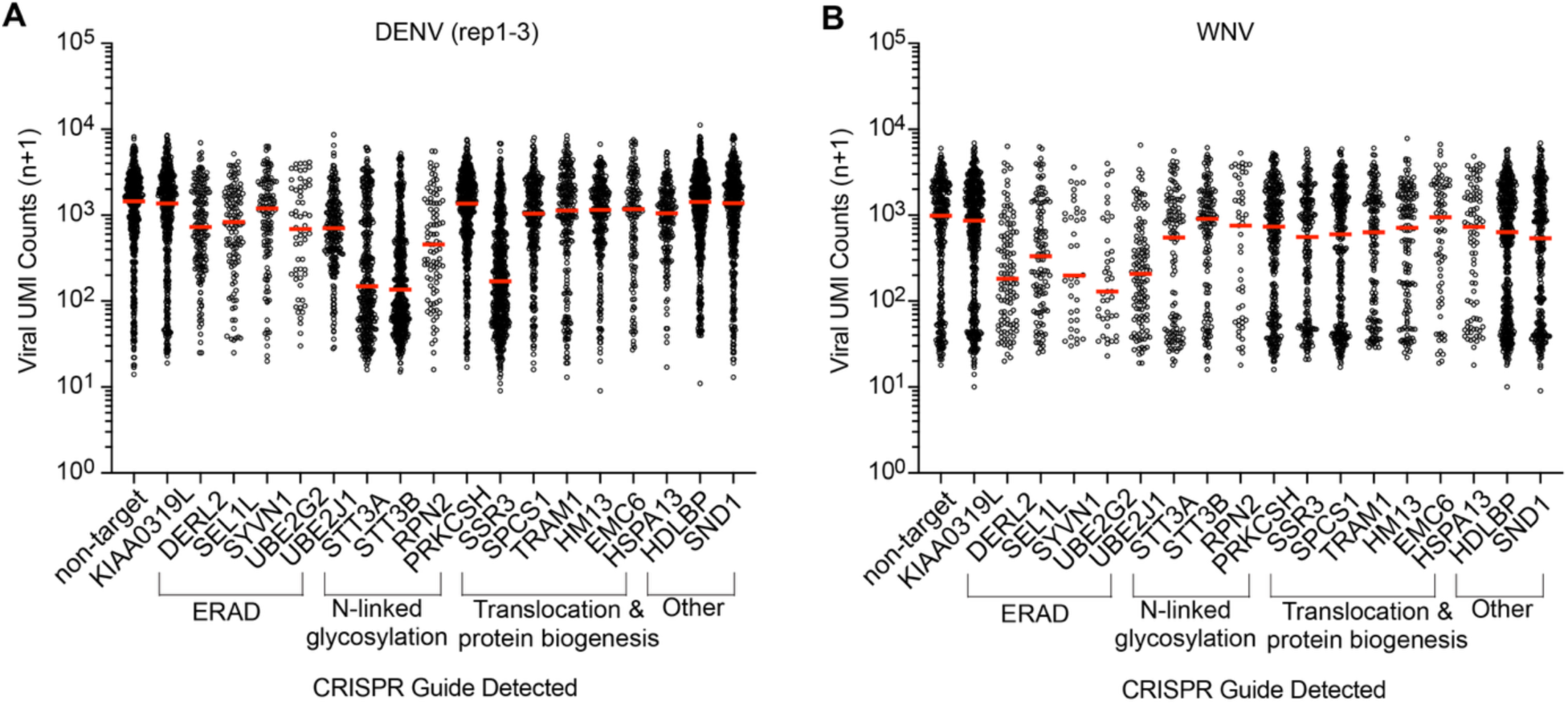
DENV and WNV QIC-seq plots. **A**, QIC-seq cell plot of DENV challenged HAP1 cells. Circles represent cells. Cells plotted by guide-detected, and DENV UMI counts (log_10_ of n+1). Red line represents median value. Replicates 1, 2 and 3 were combined. **B**, As in **A**, however HAP1 cells are challenged with WNV.

**Supplemental Figure 5:**
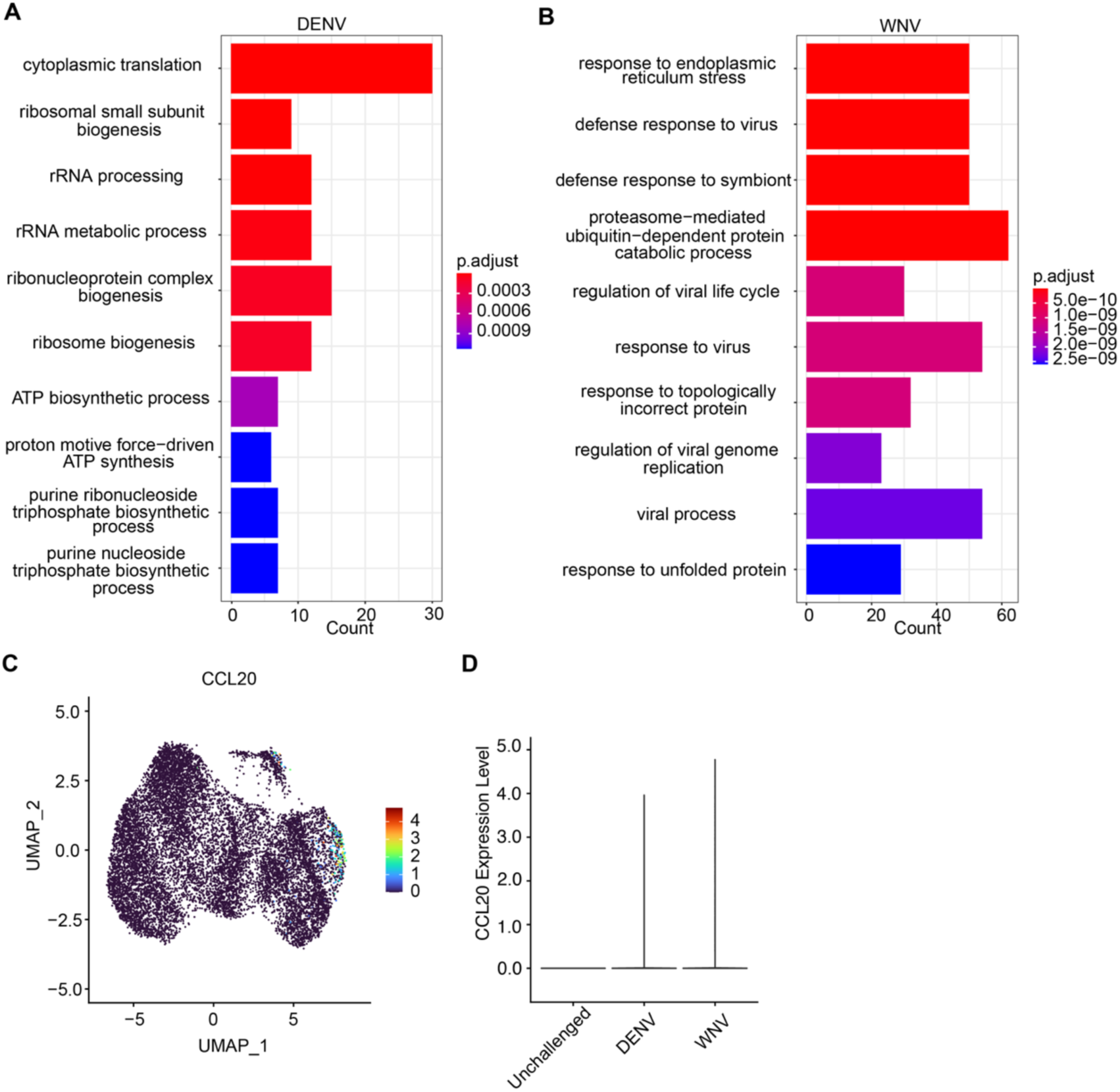
UPR and IFN response induced in WNV challenged HAP1 cells; ribosomal biogenesis induced in DENV challenged HAP1 cells. **A**, GO plot of pathways upregulated based on DEGs found between DENV challenged and unchallenged HAP1 cells. List of genes generated using log2fc 2 0.25. Top 10 most upregulated pathways plotted. Count represents the number of DEGs belonging to the indicated pathway. **B**, As in **A**, however HAP1 cells were challenged with WNV. **C**, Feature plot of all HAP1 cells, with CCL20 expression featured. **D**, Violin plot of CCL20 expression in HAP1 cells.

**Supplemental Figure 6:**
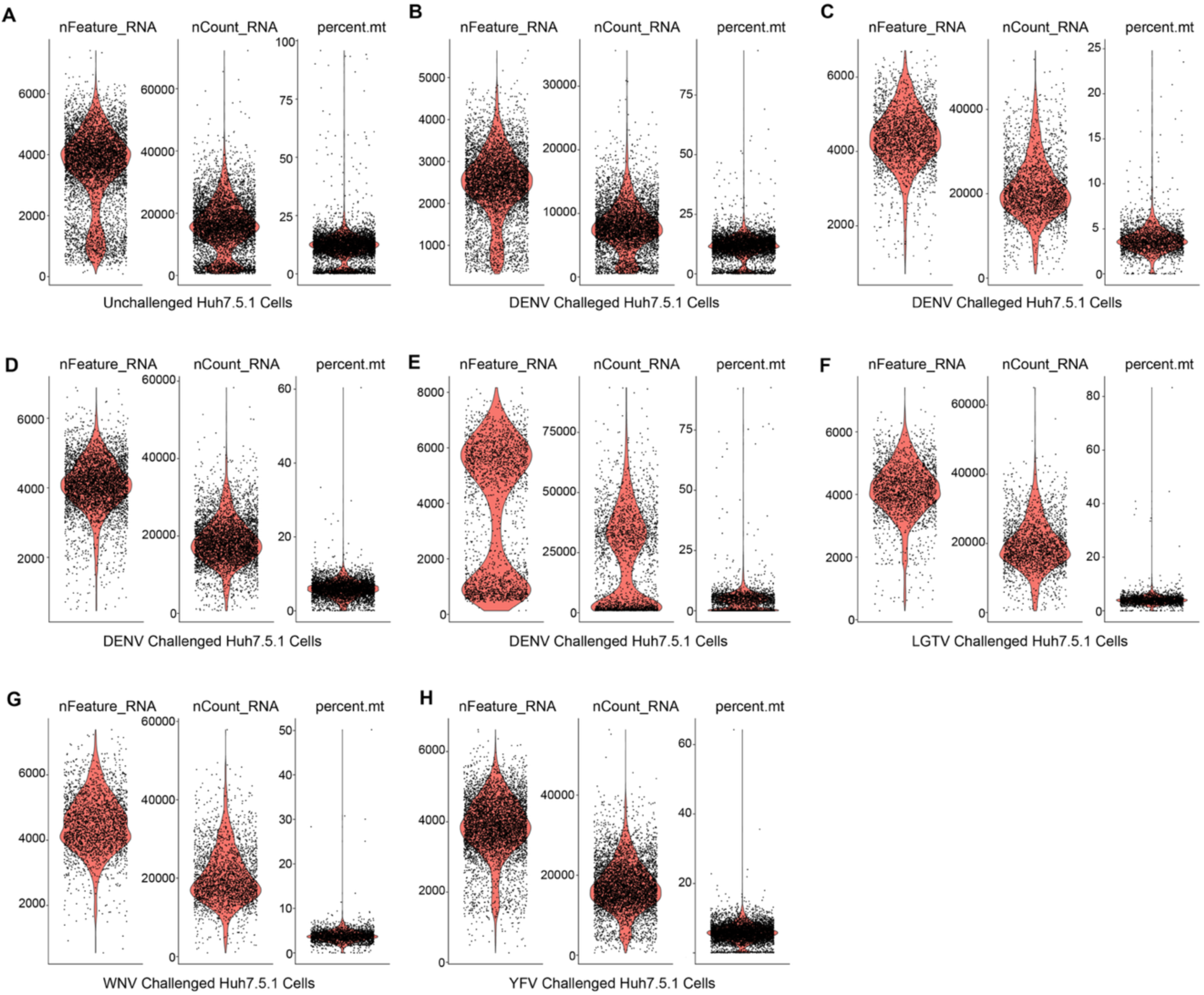
QC: Cell quality control for Huh7.5.1 cell libraries used in QIC-seq. **A-H**, Violin plots for number of RNA features, counts of RNA, and percent mitochondrial genes. Cut offs were made according to each data set (see Supplemental Table 9 for values used) before data sets were merged into final Seurat Object (SO). **A**, Unchallenged Huh7.5.1 cells. **B**, DENV-challenged Huh7.5.1 cells, biological replicate 1. **C,** DENV-challenged Huh7.5.1 cells, biological replicate 2. **D**, DENV-challenged Huh7.5.1 cells, biological replicate 3. **E**, DENV-challenged Huh7.5.1 cells, biological replicate 4. **F**, LGTV-challenged Huh7.5.1 cells. **G**, WNV-challenged Huh7.5.1 cells. **H**, YFVV-challenged Huh7.5.1 cells.

**Supplemental Figure 7:**
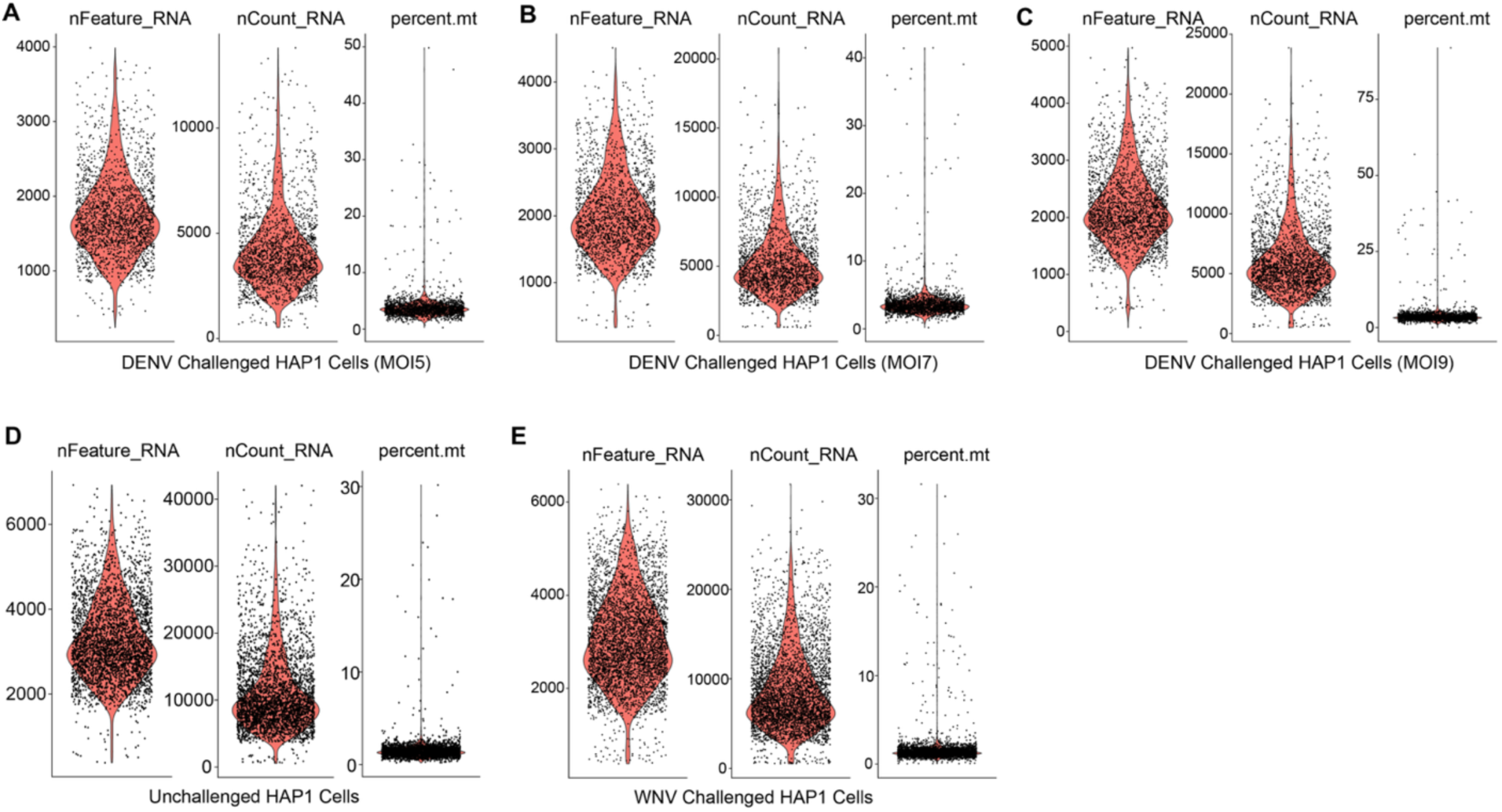
QC: Cell quality control for HAP1 cell libraries used in QIC-seq. **A-E**, Violin plots for number of RNA features, counts of RNA, and percent mitochondrial genes. Cut offs were made according to each data set (Supplemental Table 9 for values used) before data sets were merged into final Seurat Object (SO). **A**, DENV-challenged HAP1 cells (MOI5), biological replicate 1. **B**, DENV-challenged HAP1 cells (MOI7), biological replicate 2. **C**, DENV-challenged HAP1 cells (MOI9), biological replicate 3. **D**, Unchallenged HAP1 cells. **E**, WNV-challenged HAP1 cells.

## Supplementary Files

**Supplemental table 1: Virus counts feature calls by cell barcode in Huh7.5.1 cells.**

**Supplemental table 2: Gene modules.**

**Supplemental table 3: Huh7.5.1 differentially expressed genes.**

**Supplemental table 4: Virus counts and feature calls by barcode in HAP1 cells.**

**Supplemental table 5: HAP1 differentially expressed genes.**

**Supplemental table 6: Cell library oligonucleotide sequences.**

**Supplemental table 7: 10X primers and PCR conditions.**

**Supplemental table 8: Feature reference sequences.**

**Supplemental table 9: Cell quality control values.**

**Supplemental table 10: Huh7.5.1 indel primer sequences.**

## Materials and Methods

### Cell lines, generation of lentivirus and cell libraries

Huh7.5.1 cells, a kind gift from F. Chisari, were cultured in DMEM media, 10% heat-inactivated fetal bovine serum (HI-FBS), and 1% penicillin and streptomycin (pen-strep). HAP1 cells, derived from near-haploid chronic myeloid leukemia cells, KBM7, were cultured in IMDM media, 10% HI-FBS, and 1% pen-strep.

Lentivirus was generated by co-transfecting the ΔVPR, VSV-G, and pAdVAntage lentivirus packaging plasmids along with CRISPR-guide containing lentiCRISPRv2 plasmids targeting genes of interest into 293FT cells using transIT-LT1 reagent (Mirus bio). Lentivirus was collected after 48 hours, and filtered using a 0.45-mm filter. The lentiCRISPR v2 plasmid was a gift from Feng Zhang (Addgene plasmid # 52961; http://n2t.net/addgene:52961; RRID:Addgene_52961). All oligonucleotides for CRISPR sgRNA generation, QIC-seq, and indel frequency analysis ordered from Integrated DNA Technologies (IDT).

To generate the Huh7.5.1 cell library, individual knockout cell lines were generated, combined, and used for the initial QIC-seq screen. CRISPR guide RNA sequences were selected from the human GeCKO library^46^. To generate the knockout cell lines, oligos (Supplemental Table 6) were annealed, phosphorylated, and ligated into BsmBI (New England Biosciences)-cleaved lentiCRISPRv2 before transforming into Stbl3 competent bacteria and used to generate lentivirus as above. Sequencing was confirmed by Sanger sequencing. Huh7.5.1 cells were transduced with lentivirus plus 8 µg/ml protamine sulfate, and selected with 1 µg/ml puromycin after 48 hours. Cells were pooled at equal ratios. The SETD3 knockout cell line was generated as above, however was excluded from the cell library in order to represent a naïve knockout cell line in a DENV challenged environment.

To generate the HAP1 cell library, a lentivirus plasmid library was first generated, then used in a single transduction of HAP1 cells at a MOI of 0.3. To generate the HAP1 lentivirus plasmid library, 4 oligos encoding a sgRNA targeting each indicated gene and 4 non-targeting oligos were selected from the human Brunello library (Supplemental Table 6)^47^, and purchased in a pooled format. A total of 1 ng of oligo pool was amplified using the forward primer: GGCTTTATATATCTTGTGGAAAGGACGAAACACC and the reverse primer: CTAGCCTTATTTTAACTTGCTATTTCTAGCTCTAAAAC at final concentrations of 500 nM and Q5 High Fidelity polymerase (New England Biolabs) according to manufacturer’s protocol with an annealing temperature of 60°C for 20 cycles. The resulting PCR product was gel extracted, and Gibson assembled into BsmBI-cleaved lentiCRISPRv2 at a 5:1 vector to insert ratio. Assembly reaction was transformed into Endura Duo electrocompetent cells, plated to calculate gene coverage, and a small number of colonies were confirmed by Sanger sequencing.

The plasmid library was used to generate lentivirus as above. Lentivirus was collected from transfected 293FT cell supernatant, centrifuged to remove cell debris, and concentrated using PEG-it (System Biosciences). HAP1 cells were transduced with concentrated lentivirus and 8 µg/ml protamine sulfate and selected using 1 µg/ml puromycin 48 hours after transduction.

### 10X Genomics kits

All Huh7.5.1 QIC-seq screens were conducted using the 10X Genomics Chromium Next GEM Single Cell 5′ Library and Gel Bead Kit (product code 1000167), the Chromium Single Cell 5′ Library Construction Kit (product code 1000020), the Chromium Next GEM Chip G single Cell Kit (product code 1000120), and the Single Index Kit T Set A (product code 1000213). All HAP1 QIC-seq screens were conducted using 10X genomics Chromium Next GEM Single Cell 5′ Kit v2 (product code 1000265), Chromium Next GEM Chip K Single Cell Kit (product code 1000287), Chromium 5′ CRISPR kit (product number 1000451), Library Construction Kit (product number 1000190), and the Dual Index Kit TT Set A (product number 1000215). Protocols followed were Chromium Next GEM Single Cell V(D)J Reagent Kits v1.1 User Guide Rev F, and Chromium Next GEM Single Cell 5′ v2 with Feature Barcode technology_CRISPR Screening Rev B.

### Virus challenges and cell preparation for GEM generation

For Huh7.5.1 QIC-seq screens, 150,000 cells were seeded in 6 well plates in DMEM, 10% HI FBS, and 1% pen-strep, 16 hours prior to viral challenge. Cells were challenged for 48 hours with dengue virus type 2 (16681), yellow fever virus (17D), West Nile virus (Kunjin), or Langat virus (a generous gift from Marshall Bloom) all at MOI of 0.1, except Langat which was used at an MOI of 20. After 48 hours, cells were collected and resuspended in single cell suspension at 1000 cells/µl in PBS and 0.04% BSA to prevent cell aggregation. Four biological replicates were completed for DENV challenge, and single replicates were completed for Langat, West Nile and yellow fever virus QIC-seq screens. Naïve SETD3 knockout cells were added into the cell suspension of a single DENV challenge biological replicate prior to GEM generation (replicate four). HAP1 QIC-seq screens were challenged as above, however dengue was used at an MOI of 5, 7, and 9, and West Nile virus was used at an MOI of 2.5. Recovery target from GEM generation was 10,000 cells.

### QIC-seq library preparation and sequencing

For Huh7.5.1 QIC-seq screens, primers targeting the CRISPR sgRNA scaffold and the 5′ end of the indicated viral genome were added into the cell + master mix suspension at a final concentration of 250nM before loading the chromium chip (see Supplemental Table 7 for primers).

cDNA synthesis, post GEM-RT clean up, and cDNA amplification all proceeded as written in the 10X protocol. In step 3.2 of Rev F, DNA select and SPRI clean up were modified to generate two separate libraries based on DNA size, using 0.6X SPRI beads. The gene expression library (GEX) contained DNA >300 base pairs, and remained bound to beads. The viral/CRISPR library (V/C) contained DNA <300 base pairs, and was collected from the supernatant. The GEX library followed the remaining 10X genomics protocol.

The V/C library was subject to READ2 placement PCR (see Supplemental Table 7 for primers). Briefly, the SI PCR primer (10X genomics) and a primer containing the READ2 sequence was used to generate a DNA product that can be indexed for Illumina sequencing. After cDNA amplification (step 3 of rev F), and DNA selection and SPRI bead clean up (step 3.2 of rev F), 80µl of supernatant containing DNA <300 bp in length was collected. To this, 70 µl of 2X SPRIselect (Beckman) was added for a 2.0X ratio. Beads were pelleted, washed with 80% ethanol, and resuspended in 45 µl of EB buffer (Qiagen).

For READ2 placement PCR, KAPA HiFi or KAPA HiFi Hotstart polymerase (Roche) was added to 10ng of DNA from the 45 µl of bead elution. Each reaction received the 10X SI_PCR primer, and the CRISPR guide READ2 placement primer as well as the respective virus READ2 placement primers at 245 nM each (see Supplemental Table 7 for PCR conditions according to virus). Post PCR, 50 µl of 2X SPRIselect was added to the total 25 µl PCR volume for a 2.0X ratio. Beads were again pelleted, washed in 80% ethanol and resuspended in 40 µl EB buffer. From this, 5 µl were added into the indexing PCR reaction according to step 6 of rev F (see Supplemental Table 7 for indexing PCR conditions).

HAP1 QIC-seq screen modification and sequencing was completed as above, however, no primer targeting the CRISPR sgRNA scaffold was added into the cell + master mix suspension as the Chromium 5′ CRISPR kit became commercially available. All V/C and GEX libraries were prepared and indexed separately before Illumina sequencing by Novogene with an intended read target of 25 million reads per V/C library, and 50 million reads per GEX library.

### Processing of Fastq files, sgRNA indexing library (feature reference), selection of cells, and Seurat Object (SO) processing

All fastq files generated from Illumina sequencing were analyzed using 10X Genomics Cell Ranger version 4. The transcriptomic library (GEX) and viral/CRISPR (V/C) library were designated Gene Expression and CRISPR Guide Capture, respectively, in all library files. A 20 base pair sequence of the 5′ end of the viral genome, in addition to CRISPR guide sequences were included in the Feature references file. Briefly, a feature barcode sequence (BC) targeting the guide RNA scaffold (GTTTTAGAGCTAGAA) was provided, along with variable sequences specific to each guide (see Supplemental Table 8 for feature reference files). For each virus, 5′ genomic sequences were supplied as a feature, allowing for quantification from Cell Ranger’s CRISPR analysis output. Gene Expression fastq files were aligned to the Homo sapiens.GRCh38.99 genome. A final list of cells with a single type of CRISPR guide detected, meeting cell quality control standards was generated as follows and used in the QIC-seq plot and Seurat data analysis: first, Cell Ranger generated Protospacer Calls Per Cell file is used to identify a list of cells with a single type of CRISPR guide called. Next, cells present in the Filter Feature Barcode Matrix corresponding to the list of cells above is generated, and further subset according to cell quality control standards (Supplemental Table 9, Supplemental Figures 6-7). Virus counts were taken from the Filter Feature Barcode Matrix.

Data was analyzed using Seurat v.4.4.0. Seurat objects (SOs) were generated for each QIC-seq screen, and merged to generate a single SO for each cell line. For Huh7.5.1 QIC-seq screens, SOs were merged and split by type of viral challenge before undergoing SCTransformation (version 2) with cell cycle and mitochondrial mapping percentage regressed out. Lastly, integration was performed using Seurat IntegrateData() function with default parameters. To cluster and visualize Huh7.5.1 cells, principal component analysis was performed using Seurat RunPCA() function using default parameters. UMAPs were generated using the first 30 principal components as suggested by visual inspection of elbow plot, using default parameters. To identify clusters, the Seurat FindClusters() function was used with default parameters.

For HAP1 cells, all SOs were merged to a single SO, normalized, and variable features were identified using the vst selection method with 2000 features using default parameters. Data was scaled with cell cycle and mitochondrial mapping percentage regressed out. To cluster and visualize cells, Seurat RunPCA() function was used with default parameters. UMAPs were generated including the first 20 principal components as suggested by visual inspection of elbow plot, using default parameters. To identify clusters, the Seurat FindClusters() function was used with resolution 0.6 and otherwise default parameters.

### Master gene list generation

An unfolded protein response (UPR), and IFN stimulated gene (IFN) master gene list were generated and used to compare differentially expressed genes resulting from viral challenge (Supplemental Table 2). The UPR master list was generated from experimental results from Reich et al.^27^, and experimental results from Adamson et al.^26^. The IFN stimulated gene master list was generated from Shaw et al.^28^ and Lumb et al.^29^.

### QC feature scatter plots (violin plots)

All cells with a single guide detected are plotted for each QIC-seq screen. Cells with values outside indicated ranges as seen in Supplemental Table 9 are removed from analysis.

### Viral UMI count distributions

Viral UMI counts were plotted as log_10_(n+1) values using Seurat RidgePlot() function. In the Huh7.5.1 DENV QIC-seq screen, non-target guide detected cells were first subset from the processed SO. Guide detected cells were plotted, grouped by biological replicate. In the Huh7.5.1 multi-flavivirus and HAP1 QIC-seq screens, cells are grouped by viral challenge.

### QIC-Seq cell plot and normalized viral counts

Cells were grouped by detected CRISPR guide, and viral UMI counts (n+1) were plotted on log scale using GraphPad PRISM. Red line represents the median value. The QIC-seq plot for DENV challenged HAP1 cells represents three biological replicates combined. To graph normalized DENV counts, cells were grouped by CRISPR guides detected, and mean viral counts were divided by mean viral counts in non-target-1 for Huh7.5.1 guide detected cells belonging to the same QIC-seq screen or combined non-target for HAP1 guide detected cells belonging to the same QIC-seq screen. Error bars represent standard error of the mean. All four biological replicates depicted in Huh7.5.1 cells, and all three biological replicates depicted in HAP1 cells. To graph normalized viral counts in all viral challenges, cells are grouped according to viral challenge and guide detected. Mean viral UMI counts (n+1) are then divided by mean viral UMI counts of non-target-1-guide detected Huh7.5.1 cells or of combined non-target HAP1 cells belonging to the corresponding QIC-seq screen. DENV is representative of all biological replicates corresponding to cell type.

### Log1p of average RNA expression plots

All Huh7.5.1 datasets were individually subset from the final, processed SO. All virally challenged datasets were then merged with the unchallenged dataset, with the exception of the DENV challenge datasets, which were all merged together (four biological replicates), along with the unchallenged dataset. The log1p average RNA expression was calculated for each gene in both data sets, and plotted as a single circle. The y=x line shown indicates equal gene expression between challenged and unchallenged cells. From the RNA assay, the Seurat FindMarkers() function was used to identify differentially expressed genes between the virally challenged and unchallenged cells. Differentially expressed genes identified with a log2fc value ≥ ±0.25 were cross referenced with the UPR, and IFN stimulated master gene lists, and plotted as red, or dark green circles, respectively. Genes corresponding to the UPR and IFN stimulated master lists that were not differentially expressed were plotted as pink and light green, respectively.

### Module scores

A module score (a score given to a cell that is indicative of how upregulated a specific list of genes is, relative to other random genes in a dataset) was assigned to each cell in the original processed Seurat Object based on the UPR and IFN stimulated master gene list (Supplemental Table 2) using the Seurat function AddModuleScore(). All module scores were assigned prior to any subsetting and further analysis. The Seurat VlnPlot() function was used to plot the module score according to specific parameters, such as by viral challenge, by guide, or by type of perturbation.

### GO analysis

Genes were identified as differentially expressed using the Seurat FindMarkers() function, with a log2fc threshold of 0.1, using the default Wilcoxon Rank Sum test. Non-mitochondrial genes were then further subset using a cut off of log2fc ≥ ±0.25, and adjusted p value ≥ 0.05. The resulting differentially expressed genes were then used for GO analysis using the enrichGO() function from clusteprofiler48, with the top 10 most upregulated pathways plotted.

### Indel frequency

Genomic DNA from approximately 8 million cells from each Huh7.5.1 knockout cell line was extracted using the Qiagen DNeasy Blood and Tissue kit. 145 ng of DNA of all but three knockout cell lines was subject to PCR in which a 200-280 base pair length of genomic DNA encoding the target site was amplified (see Supplemental Table 10 for primers). In the case of the DERL2, SND1, and TRAM1 knockout cell lines, DNA was subject to PCR in which a 400-600 base pair length of genomic DNA encoding the target site was amplified. Genomic DNA encoding the sgRNA target site was amplified using the Q5 High Fidelity polymerase, and underwent cycling conditions according to manufacturer’s protocol, with an anneal temperature of 65°C for 40 cycles. Samples were sequenced by the Massachusetts General Hospital Center for Computational and Integrative Biology for CRISPR or complete amplicon sequencing. CRISPR sequencing results were analyzed using publicly available CRISPResso2^49^. DERL2, SND1, and TRAM1 sequencing was returned as consensus sequences, and manually analyzed for likely loss-of-function.

### Dimensional reduction analysis and feature visualization

All feature plots were generated using Seurat’s FeaturePlot() function. To visualize viral counts, module scores, or indicated gene expression in cells the RNA assay was used.

### Correlations

All data sets were subset by viral challenge, and Spearman’s correlation coefficient was calculated between viral UMI counts and module scores in R. Correlations were then plotted using corrplot^50^.

## Acknowledgements

We thank members of the Carette lab including Michael Zobitz Palo and Alejandro Matia for thoughtful feedback and discussion on experimental design, data processing and figure presentation. We thank Jessica Fessler for assistance with R scripts, Daphne Cooper and Christine Kao at 10X Genomics for protocol discussions, and the Stanford Genetics Core for assistance with 10X protocol design and execution, and use of the BSL2 Chromium Controller. We thank the Kopito lab for project feedback. We thank Pricilla Yang for feedback and kind mentorship. We thank Michael J Bennett for his assistance with initial Fastq alignments, and for his unwavering support. Protocol design in Figs. 1a, Supplemental Fig. 1e, and 4a were generated using Biorender. This work was funded by the National Institutes of Health (NIH) R01 AI169467 (to J.E.C), NIH R01 AI140186 (to J.E.C) , NIH R01 AI141970 (to J.E.C), NIH T32 AI732834 (to A.J.D. and B.S.W.), Burroughs Wellcome Fund of Investigators in the Pathogenesis of Infectious Disease (to J.E.C.) Stanford University School of Medicine Dean’s Postdoctoral fellowship (to B.S.W.), and A.P. Giannini Postdoctoral Research Fellowship (to B.S.W.).

## Contributions

J.E.C., A.J.D., B.S.W, and J.Z. contributed to project conception. J.E.C., A.J.D., and B.S.W. contributed to experimental design. J.Z. generated the Huh7.5.1 cell library. A.J.D generated the HAP1 cell library and performed QIC-seq screens. A.J.D. wrote scripts used to analyze data, A.J.D. and B.S.W. analyzed data. J.E.C., A.J.D., and B.S.W. wrote the manuscript. J.E.C. supervised the research. All of the authors read and approved the final version of this manuscript before submission.

## References

1. Bhatt, S. et al. The global distribution and burden of dengue. Nature 496, 504–507 (2013).

2. Pierson, T. C. & Diamond, M. S. The continued threat of emerging flaviviruses. Nat. Microbiol. 5, 796–812 (2020).

3. Dobler, G., Gniel, D., Petermann, R. & Pfeffer, M. Epidemiology and distribution of tick-borne encephalitis. Wien. Med. Wochenschr. 162, 230–238 (2012).

4. Kwasnik, M., Rola, J. & Rozek, W. Tick-borne encephalitis-review of the current status. J. Clin. Med. 12, (2023).

5. Hills, S. L., Poehling, K. A., Chen, W. H. & Staples, J. E. Tick-borne encephalitis vaccine: Recommendations of the Advisory Committee on Immunization Practices, United States, 2023. MMWR Recomm. Rep. 72, 1–29 (2023).

6. Tabachnick, W. J. Climate change and the arboviruses: Lessons from the evolution of the dengue and yellow fever viruses. Annu. Rev. Virol. 3, 125–145 (2016).

7. Barrows, N. J. et al. Biochemistry and molecular biology of flaviviruses. Chem. Rev. 118, 4448–4482 (2018).

8. Paul, D. & Bartenschlager, R. Flaviviridae replication organelles: Oh, what a tangled web we weave. Annu. Rev. Virol. 2, 289–310 (2015).

9. Krishnan, M. N. et al. RNA interference screen for human genes associated with West Nile virus infection. Nature 455, 242–245 (2008).

10. Ma, H. et al. A CRISPR-based screen identifies genes essential for west-Nile-virus-induced cell death. Cell Rep. 12, 673–683 (2015).

11. Marceau, C. D. et al. Genetic dissection of Flaviviridae host factors through genome-scale CRISPR screens. Nature 535, 159–163 (2016).

12. Lin, D. L. et al. Dengue virus hijacks a noncanonical oxidoreductase function of a cellular oligosaccharyltransferase complex. MBio 8, (2017).

13. Savidis, G. et al. Identification of Zika virus and dengue virus dependency factors using functional genomics. Cell Rep. 16, 232–246 (2016).

14. Shah, P. S. et al. Comparative Flavivirus-host protein interaction mapping reveals mechanisms of dengue and Zika virus pathogenesis. Cell 175, 1931–1945.e18 (2018).

15. Zhang, R. et al. A CRISPR screen defines a signal peptide processing pathway required by flaviviruses. Nature 535, 164–168 (2016).

16. Peña, J. & Harris, E. Dengue virus modulates the unfolded protein response in a time-dependent manner. J. Biol. Chem. 286, 14226–14236 (2011).

17. Lewy, T. G., Grabowski, J. M. & Bloom, M. E. BiP: Master regulator of the unfolded protein response and crucial factor in Flavivirus biology. Yale J. Biol. Med. 90, 291–300 (2017).

18. Lewy, T. G. et al. PERK-mediated unfolded protein response signaling restricts replication of the tick-borne Flavivirus Langat virus. Viruses 12, 328 (2020).

19. Ooi, Y. S. et al. An RNA-centric dissection of host complexes controlling flavivirus infection. Nat. Microbiol. 4, 2369–2382 (2019).

20. Blight, K. J., McKeating, J. A. & Rice, C. M. Highly permissive cell lines for subgenomic and genomic hepatitis C virus RNA replication. J. Virol. 76, 13001–13014 (2002).

21. Balsitis, S. J. et al. Tropism of dengue virus in mice and humans defined by viral nonstructural protein 3-specific immunostaining. Am. J. Trop. Med. Hyg. 80, 416–424 (2009).

22. Mimitou, E. P. et al. Multiplexed detection of proteins, transcriptomes, clonotypes and CRISPR perturbations in single cells. Nat. Methods 16, 409–412 (2019).

23. Gaunt, M. W. et al. Phylogenetic relationships of flaviviruses correlate with their epidemiology, disease association and biogeography. J. Gen. Virol. 82, 1867–1876 (2001).

24. Pleiner, T. et al. Structural basis for membrane insertion by the human ER membrane protein complex. Science 369, 433–436 (2020).

25. Verhaegen, M. & Vermeire, K. The endoplasmic reticulum (ER): a crucial cellular hub in flavivirus infection and potential target site for antiviral interventions. Npj Viruses 2, 24 (2024).

26. Adamson, B. et al. A multiplexed single-cell CRISPR screening platform enables systematic dissection of the unfolded protein response. Cell 167, 1867–1882.e21 (2016).

27. Reich, S. et al. A multi-omics analysis reveals the unfolded protein response regulon and stress-induced resistance to folate-based antimetabolites. Nat. Commun. 11, 2936 (2020).

28. Shaw, A. E. et al. Fundamental properties of the mammalian innate immune system revealed by multispecies comparison of type I interferon responses. PLoS Biol. 15, e2004086 (2017).

29. Lumb, J. H. et al. DDX6 represses aberrant activation of interferon-stimulated genes. Cell Rep. 20, 819–831 (2017).

30. Labeau, A. et al. A genome-wide CRISPR-Cas9 screen identifies the dolichol-phosphate mannose synthase complex as a host dependency factor for dengue virus infection. J. Virol. 94, (2020).

31. Zanini, F. et al. Virus-inclusive single-cell RNA sequencing reveals the molecular signature of progression to severe dengue. Proc. Natl. Acad. Sci. U. S. A. 115, E12363–E12369 (2018).

32. Zanini, F., Pu, S.-Y., Bekerman, E., Einav, S. & Quake, S. R. Single-cell transcriptional dynamics of flavivirus infection. Elife 7, (2018).

33. Russell, A. B., Elshina, E., Kowalsky, J. R., te Velthuis, A. J. W. & Bloom, J. D. Single-cell virus sequencing of influenza infections that trigger innate immunity. J. Virol. 93, (2019).

34. Russell, A. B., Trapnell, C. & Bloom, J. D. Extreme heterogeneity of influenza virus infection in single cells. Elife 7, (2018).

35. Sunshine, S. et al. Systematic functional interrogation of SARS-CoV-2 host factors using Perturb-seq. Nat. Commun. 14, 6245 (2023).

36. Kaufmann, S. H. E., Dorhoi, A., Hotchkiss, R. S. & Bartenschlager, R. Host-directed therapies for bacterial and viral infections. Nat. Rev. Drug Discov. 17, 35–56 (2018).

37. Colpitts, T. M., Barthel, S., Wang, P. & Fikrig, E. Dengue virus capsid protein binds core histones and inhibits nucleosome formation in human liver cells. PLoS One 6, e24365 (2011).

38. Zhang, K. et al. Endoplasmic reticulum stress activates cleavage of CREBH to induce a systemic inflammatory response. Cell 124, 587–599 (2006).

39. Vabret, N. et al. Y RNAs are conserved endogenous RIG-I ligands across RNA virus infection and are targeted by HIV-1. iScience 25, 104599 (2022).

40. Sumpter, R., Jr et al. Regulating intracellular antiviral defense and permissiveness to hepatitis C virus RNA replication through a cellular RNA helicase, RIG-I. J. Virol. 79, 2689– 2699 (2005).

41. Guo, J.-T., Hayashi, J. & Seeger, C. West Nile virus inhibits the signal transduction pathway of alpha interferon. J. Virol. 79, 1343–1350 (2005).

42. Liu, W. J. et al. Inhibition of interferon signaling by the New York 99 strain and Kunjin subtype of West Nile virus involves blockage of STAT1 and STAT2 activation by nonstructural proteins. J. Virol. 79, 1934–1942 (2005).

43. Espada-Murao, L. A. & Morita, K. Delayed cytosolic exposure of Japanese encephalitis virus double-stranded RNA impedes interferon activation and enhances viral dissemination in porcine cells. J. Virol. 85, 6736–6749 (2011).

44. Uchida, L. et al. The dengue virus conceals double-stranded RNA in the intracellular membrane to escape from an interferon response. Sci. Rep. 4, 7395 (2014).

45. Aguirre, S. et al. Dengue virus NS2B protein targets cGAS for degradation and prevents mitochondrial DNA sensing during infection. Nat. Microbiol. 2, (2017).

46. Shalem, O. et al. Genome-scale CRISPR-Cas9 knockout screening in human cells. Science 343, 84–87 (2014).

47. Doench, J. G. et al. Optimized sgRNA design to maximize activity and minimize off-target effects of CRISPR-Cas9. Nat. Biotechnol. 34, 184–191 (2016).

48. Yu, G. Thirteen years of clusterProfiler. Innovation (Camb.) 5, 100722 (2024).

49. Clement, K. et al. CRISPResso2 provides accurate and rapid genome editing sequence analysis. Nat. Biotechnol. 37, 224–226 (2019).

50. Wei, T. and Simko, V. (2017) R Package “Corrplot”: Visualization of a Correlation Matrix (Version 0.84). https://github.com/taiyun/corrplot.

